# High-resolution dissection of human cell type-specific enhancers in *cis* and *trans* activities

**DOI:** 10.1101/2023.09.23.559140

**Authors:** Meng Wang, Xiaoxu Yang, Qixi Wu

## Abstract

The spatiotemporal specific gene expression is regulated by cell type-specific regulatory elements including enhancers, silencers and insulators etc. The massively parallel reporter assay (MPRA) methods like STARR-seq facilitate the systematic study of DNA sequence intrinsic enhancer activities in a large scale. However, when applied to human cells, it remains challenging to identify and quantify cell type-specific active enhancers in the genome-wide scale with high-resolution, due to the large size of human genome. In this study, we selected the H3K4me1 associated dinucleosome with the linker DNA sequences as candidate enhancer sequences in two different human cell lines and performed ChIP-STARR-seq to quantify the cell type-specific enhancer activities with high-resolution in a genome-wide scale. Furthermore, we investigated how the activity landscape of enhancer repository would change when transferred from native cells (*cis* activity) to another cell lines (*trans* activity). Using ChIP-STARR-seq of the candidate enhancers in native cells and another type of cells, we obtained enhancers *cis* activity maps and *trans* activity maps in two different cell lines. The *cis* and *trans* activity maps enabled us to identify cell type-specific active enhancers, with enrichment of motifs of differentially expressed TFs. Comparisons between the *cis* and *trans* activity maps revealed general consistent regulatory property with different levels of activity in the two cell types, suggesting the sequence intrinsic regulatory properties keep similar in different type of cells. This study provides a new perspective of sequence intrinsic enhancer activities in different types of cells.

## Introduction

The cell type-specific gene expression program is orchestrated by the regulatory elements across the genome and the specific repository of transcription factors (TFs) binding to these elements (Gasperini et al. 2020; Rao et al. 2021). Genome-wide systematic identification of regulatory elements like enhancers can be achieved by sequencing the defined combination of histone modification markers, chromatin accessibilities, or capturing the bidirectional enhancer RNAs (eRNA) (Shlyueva et al. 2014). In addition, the massively parallel reporter assay (MPRA) has been widely employed to high-throughput directly quantify the intrinsic enhancer activity of DNA sequences out of the native chromatin state (Patwardhan et al. 2012; Ernst et al. 2016; Klein et al. 2020; Sahu et al. 2022; Zhao et al. 2023). STARR-seq (Arnold et al. 2013) is one of the popular MPRA method that puts the test DNA sequence downstream of the reporter gene while upstream of the poly-adenylation signal sequence, which enables the transcribed reporter RNAs containing the test sequence itself. Thus, by high-throughput sequencing of the reporter RNA and mapping the test sequence back to genome, it can not only identify the genomic location of enhancer sequences, but also provide a quantitative map of enhancer activities across the genome.

STARR-seq has been widely employed in genome-wide quantification of enhancer activities in different species and different types of cells (Das et al. 2023). When applied to human and other mammalian cells (Liu et al. 2017; Johnson et al. 2018), due to the large size of their genomes, usually large fragment sizes have to be adopted if the whole genome is used as input, resulting in low-resolution enhancer sequences identified. Fortunately, candidate enhancers across the genome can be narrowed down to regions with specific histone modifications like H3K27ac and H3K4me1, or open chromatin regions, or specific TF binding regions (Gasperini et al. 2020). To obtain high-resolution enhancer activity maps in human and other mammalian cells, methods combining the capture of a subset of the genome together with STARR-seq have been developed including CapSTARR-seq (Vanhille et al. 2015), ChIP-STARR-seq (Vockley et al. 2016; Barakat et al. 2018), ATAC-STARR-seq (Wang et al. 2018b; Hansen and Hodges 2022) and FAIRE-STARR-seq (Glaser et al. 2021). Studies using these methods have revealed diverse cell type-specific enhancer activities in different conditions. However, current studies mainly investigate enhancer activities in the native cells (*cis* activity) (Yanez-Cuna et al. 2014).

Here, we asked what the enhancer activities would be when the same set of candidate enhancer sequences are transfected to the native cells (*cis* activity), and when they are transfect to another different type of cells (*trans* activity). The *trans* activity of enhancers is only studied in a small limited scale, lacking genome-wide systematic studies (Mattioli et al. 2020). To investigate the genome-wide enhancers *cis* activities and *trans* activities in high-resolution, we performed ChIP-STARR-seq in two different human cell lines K562 and A549, by isolating the H3K4me1 associated DNA as candidate enhancers in each cell line, transfected the two sets of candidate enhancer sequences to their native cells and cross-transfected each set to the other cell line (Fig. 1A). We identified cell type-specific active enhancers, with different TFs motif enrichment, corresponding to the differential expression of these TFs in the two types of cells. Comparisons between the *trans* activities and *cis* activities of the same set of DNA sequences revealed general consistent regulatory property with different levels of activity in the two cell lines, suggesting for the same DNA sequence, its intrinsic regulatory property keeps similar in different types of cells.

**Figure 1.**
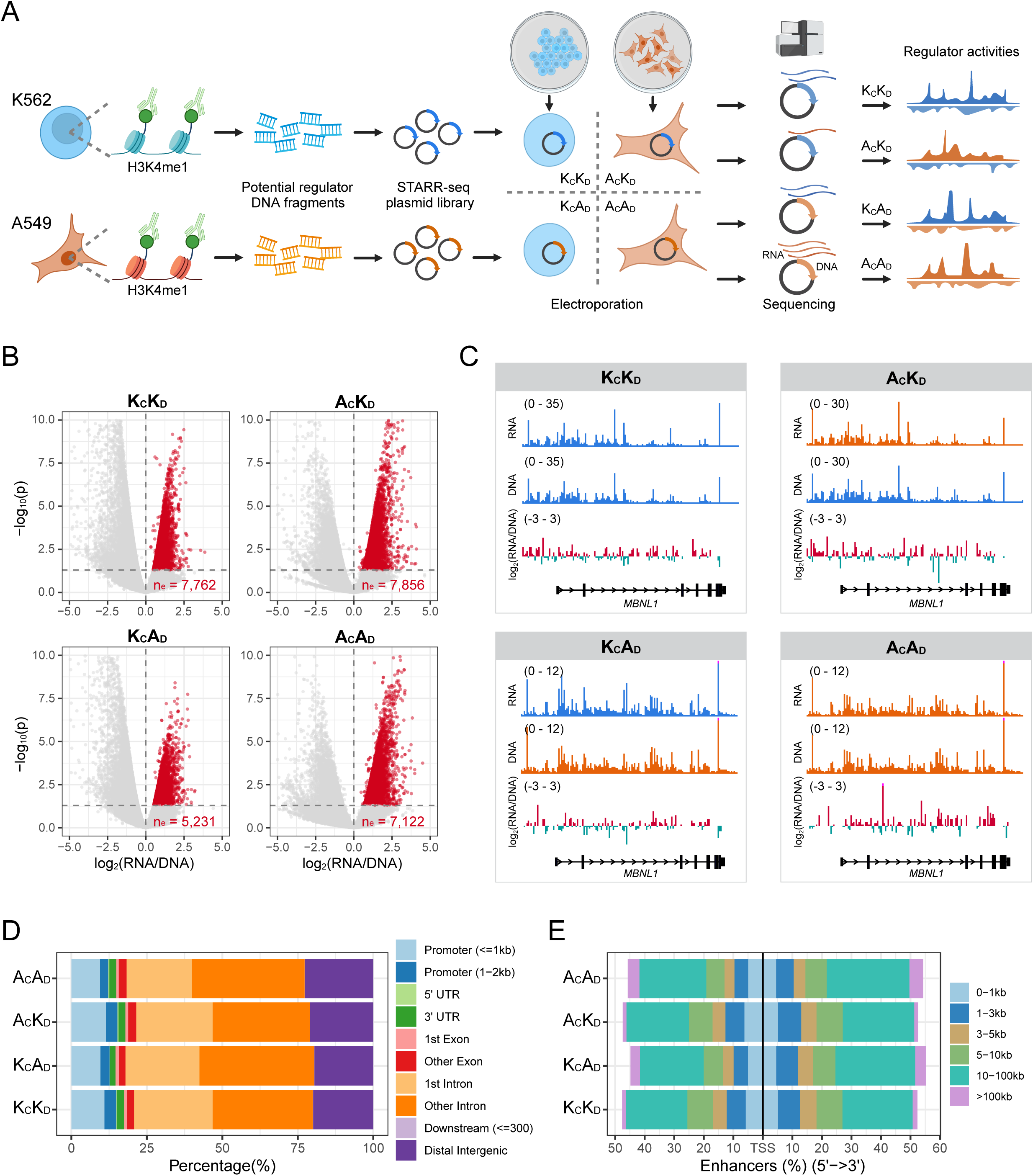
Enhancer activity maps in *cis* and *trans* for K562 and A549 cells generated by H3K4me1 ChIP-STARR-seq. (A) Experimental design and procedures of ChIP-STARR-seq in *cis* and *trans* condition. K_C_K_D_ – K562 cells with K562 DNA; A_C_A_D_ – A549 cells with A549 DNA; A_C_K_D_ – A549 cells with K562 DNA; K_C_A_D_ – K562 cells with A549 DNA. (B) Volcano plot showing the identified active enhancers in each condition. (C) Example genome browser snapshot showing the reporter RNA FPKM signals, isolated plasmid DNA FPKM signals and the calculated enhancer activities measured by the log2 fold change between RNA signals and DNA signals. (D) Genomic distribution of the active enhancers identified in each condition. (E) Distance distribution of the active enhancers to their nearest transcription start site (TSS) in each condition.

## Results

### Genome-wide high-resolution enhancer activity maps in *cis* and *trans* by ChIP-STARR-seq

To systematically study the enhancers activities in *cis* and *trans*, we choose two different human cell lines, the K562 cell line which is a bone marrow derived myelogenous leukemia cell line, and the A549 cell line which is an epithelial cell line derived from human lung adenocarcinoma. We performed chromatin immunoprecipitation (ChIP) of H3K4me1 to capture the DNA fragments associated with this histone marker and used them as the input for the following STARR-seq to screen active enhancers. We used H3K4me1 associated DNA as candidate enhancers because H3K4me1 marks both active enhancers and poised enhancers, while H3K27ac only marks active enhancers (Calo and Wysocka 2013). This enabled us to study all the potential enhancers in the genome-wide scale. To achieve high-resolution dissection of enhancer activities, we captured the dinucleosomes with linker DNA between them, resulting in about 350bp DNA fragments, by controlling the concentration and reaction time of MNase digestion to enrich dinucleosomes followed by precise DNA size selection (Supplemental Fig. S1A). We reasoned that TFs (except pioneer factors) would bind to the linker DNA between two successive nucleosomes so the H3K4me1 marked dinucleosome together with the linker DNA would provide the high-resolution fine-scale candidate enhancer sequences.

For each of the two cell lines, we performed two replicates of H3K4me1 ChIP. We first evaluated these ChIP fragments using high-throughput sequencing (ChIP-seq). The replicates in each type of cells showed high consistency with Pearson correlation coefficient 0.95 in K562 and 0.92 in A549 (Supplemental Fig. S1B). We obtained 72,764 and 73,196 reproducible H3K4me1 peaks in K562 and A549, respectively. The H3K4me1 peak regions showed the enrichment of H3K27ac, high chromatin accessibility and p300 binding in both cell lines (Supplemental Fig. S1C,D), which were consistent with the known common features of enhancers (Calo and Wysocka 2013). Most of the H3K4me1 regions were cell type-specific (Supplemental Fig. S1E), showing highly diverse enhancer repositories in different types of cells. For both cell lines, around 70% of H3K4me1 peaks were in intronic regions or distal genomic regions (Supplemental Fig. S1F), which were far away from gene transcription start sites (TSSs) (Supplemental Fig. S1G).

Next, we used the H3K4me1 ChIP DNA fragments as input to construct and amplified the STARR-seq plasmid library (ChIP-STARR-seq) with the plasmid origin of replication (ORI) as the core promoter (Muerdter et al. 2018). Each plasmid library was transfected to the corresponding native cells and cross-transfected to the other cell line by electroporation (Fig. 1A). This enabled us to obtain four activity maps of enhancers – two *cis* activity maps (K_C_K_D_ – K562 cells with K562 DNA; A_C_A_D_ – A549 cells with A549 DNA), and two *trans* activity maps (A_C_K_D_ – A549 cells with K562 DNA; K_C_A_D_ – K562 cells with A549 DNA). We evaluated the electroporation efficiency by measuring the percentage of GFP+ cells. The efficiency was as high as ∼97% for K562 cells and for A549 cells the electroporation efficiency was ∼50% (Supplemental Fig. S2A). After electroporation and culturing for 24h, we extracted both reporter RNAs and the transfected plasmid DNAs from the cells for sequencing. For each condition, we performed two replicates (Supplemental Fig. S2B,C). To reliably and accurately quantify candidate enhancer activities (Hansen and Hodges 2022), we used the plasmid DNA as control and leveraged the two replicates to calculate the p-value and the fold change of reporter RNA over plasmid DNA by DESeq2 (Love et al. 2014) (see Methods). The significant regions (p-value < 0.05) with positive log2 RNA over DNA fold change were identified as enhancers in each condition (Fig. 1B). Finally, for the *cis* activities, we obtained 7,762 enhancers in K562 cells (K_C_K_D_) and 7,122 enhancers in A549 cells (A_C_A_D_). For the *trans* activities, we obtained 7,856 enhancers in A549 cells transfected with K562 DNA (A_C_K_D_), and 5,231 enhancers in K562 cells transfected with A549 DNA (K_C_A_D_). Figure 1C provided examples of the activities map in each condition, which illustrated diverse activity patterns in different conditions for the same genomic region. The function of the nearest genes of the active enhancers in K562 were enriched in protein polyubiquitination, histone modification and RAS signal transduction etc., while the function of the nearest genes of A549 active enhancers were enriched in cell polarity, actin filament organization and response to transforming growth factor beta etc. (Supplemental Fig. S2D). More than 75% of the active enhancers were located in intron or distal regions (Fig. 1D) with 5 – 100kb away from nearest transcription start sites (Fig. 1E), while ∼15% of the active enhancers were in promoter regions, which was in line with the findings that promoter sequences can also function as enhancers (Nguyen et al. 2016).

### Cell type-specific enhancers show enrichment of motifs of differentially expressed TFs

We investigated the *cis* activity maps first to study the cell type-specific enhancers in K562 and A549 cells. We investigated the chromatin features in these active enhancer regions in each cell line. For both cell lines, compared to the negative sequences, the active enhancer sequences showed higher H3K27ac enrichment, chromatin opening and p300 binding (Fig. 2A and Supplemental Fig. S3A), indicting most of them were also active in their native chromatin environment. Indeed, when compared to the ChromHMM annotation (Ernst and Kellis 2012) which is based on the chromatin state, most of the active enhancers were overlapped with ChromHMM enhancers annotation (Fig. 2B), while the negative sequences were not (Supplemental Fig. S3B).

**Figure 2.**
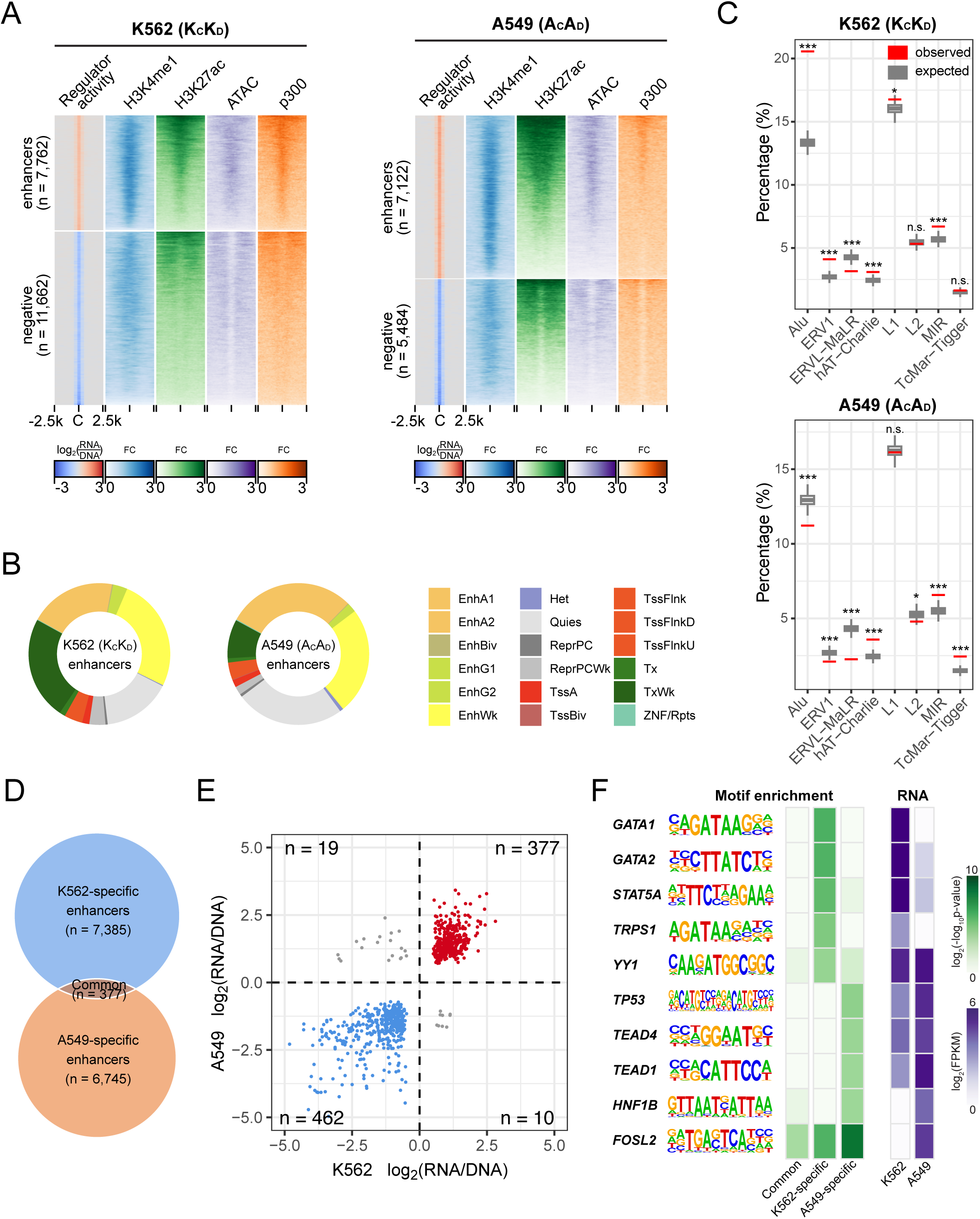
Cell type-specific active enhancers in K562 and A549. (A) Heatmaps showing the enrichment of H3K4me1, H3K27ac, chromatin accessibility (measured by ATAC-seq) and p300 binding signal at the genomic regions of active enhancers compared to negative sequences in K562 cells (left) and in A549 cells (right). (B) Overlaps of active enhancers with ChromHMM annotation. EnhA1 - Active enhancer 1; EnhA2 - Active enhancer 2; EnhBiv - Bivalent enhancer; EnhG1 - Genic enhancer 1; EnhG2 - Genic enhancer 2; EnhWk - Weak enhancer; Het - Heterochromatin; Quies - Quiescent/low; ReprPC - Repressed Polycomb; ReprPCWk - Weak repressed Polycomb; TssA - Active TSS; TssBiv - Bivalent/poised TSS; TssFlnk - Flanking TSS; TssFlnkD - Flanking TSS downstream; TssFlnkU - Flanking TSS upstream; Tx - Strong transcription; TxWk - Weak transcription; ZNF/Rpts - ZNF genes & repeats. (C) Overlaps of active enhancers with transposon elements. *: p-value<0.05, **: p-value<0.01, ***: p- value<0.001, n.s.: not significant. (D) Overlaps between K562 active enhancers and A549 active enhancers. (E) Activities comparison of the common significant active enhancers and common significant negative sequences between K562 and A549. (F) The enriched motifs of TFs in each group of active enhancers and the expression levels of the TFs in K562 and A549.

It has been reported that functional enhancers could evolve from transposon elements which are prevalent across the genome (Judd et al. 2021; Modzelewski et al. 2022). We analyzed whether specific types of transposons were overrepresented or underrepresented in the active enhancers of each type of cells. Results showed that hAT and MIR transposons were overrepresented in the active enhancers of both K562 and A549 cells (Fig. 2C). The K562 active enhancers had significantly higher than expected overlaps with the Alu and ERV1 transposons. However, these two types of transposons were significantly underrepresented in A549 active enhancers. These results suggested different types of transposon elements had different contributions to the functional enhancers in different types of cells.

The comparison between the active enhancers in K562 and that in A549 showed most of them were cell type-specific, with only 377 overlapped enhancers (Fig. 2D), while 7,385 enhancers were specific to K562, and 6,745 enhancers were specific to A549. For the overlapped enhancers and the overlapped negative sequences, they generally kept the same regulatory property in the two cell lines but with different activity levels (Fig. 2E and Supplemental Fig. S3C). We compared the TFs motif enrichment between these cell type-specific enhancers and found the cell type-specific enriched TFs that showed differential expression in the two cell lines (Fig. 2F). Motifs of GATA factors *GATA1/2*, *STAT5A* and *TRPS1* were specifically enriched in the K562- specific enhancers, and these TFs were only highly expressed in K562 cells. For A549-specific enhancers, *HNF1B* motif were specifically enriched and only highly expressed in A549 cells. Motifs of *TP53* and the Hippo pathway key factors *TEAD1/4* were specifically enriched in A549 enhancers, but these factors were highly expressed in both cell lines. For the cell type-specific negative sequences, we observed the enrichment of repressor TF motifs including *GFI1B* and *BACH1* in K562 cells, and *E2F3*, *E2F6* and *CTCF* in both K562 and A549 cells (Supplemental Fig. S3D). These results showed the high diversity of cell type-specific enhancers with different TFs binding potential.

### Enhancer activities in *trans* reveal similar regulatory properties with different activity levels

Next, utilizing the *trans* activity maps, we investigated how the enhancer activities would change when the enhancer repository from one type of cells was transferred into another type of cells. We found that overall, the activities in *trans* were generally consistent with that in the native cells (Fig. 3A,B), with Spearman correlation coefficient 0.64 and 0.53 for K562 DNA and A549 DNA, respectively. When only considering the significant enhancers and negative sequences in both cell lines, the trend kept similar (Fig. 3C,D): the enhancers in *cis* condition tended to be also enhancers in *trans* condition, while the negative sequences in *cis* condition tended to be also negative in *trans* condition. Although the enhancer and repressor properties kept unchanged between *cis* and *trans* conditions, the levels of activity were quite different between *cis* and *trans* conditions (Fig. 3E,F). While a portion of enhancers had high activities in both *cis* and *trans*, most enhancers preferred to show higher activities in either *cis* or *trans* condition (Fig. 3E,F).

**Figure 3.**
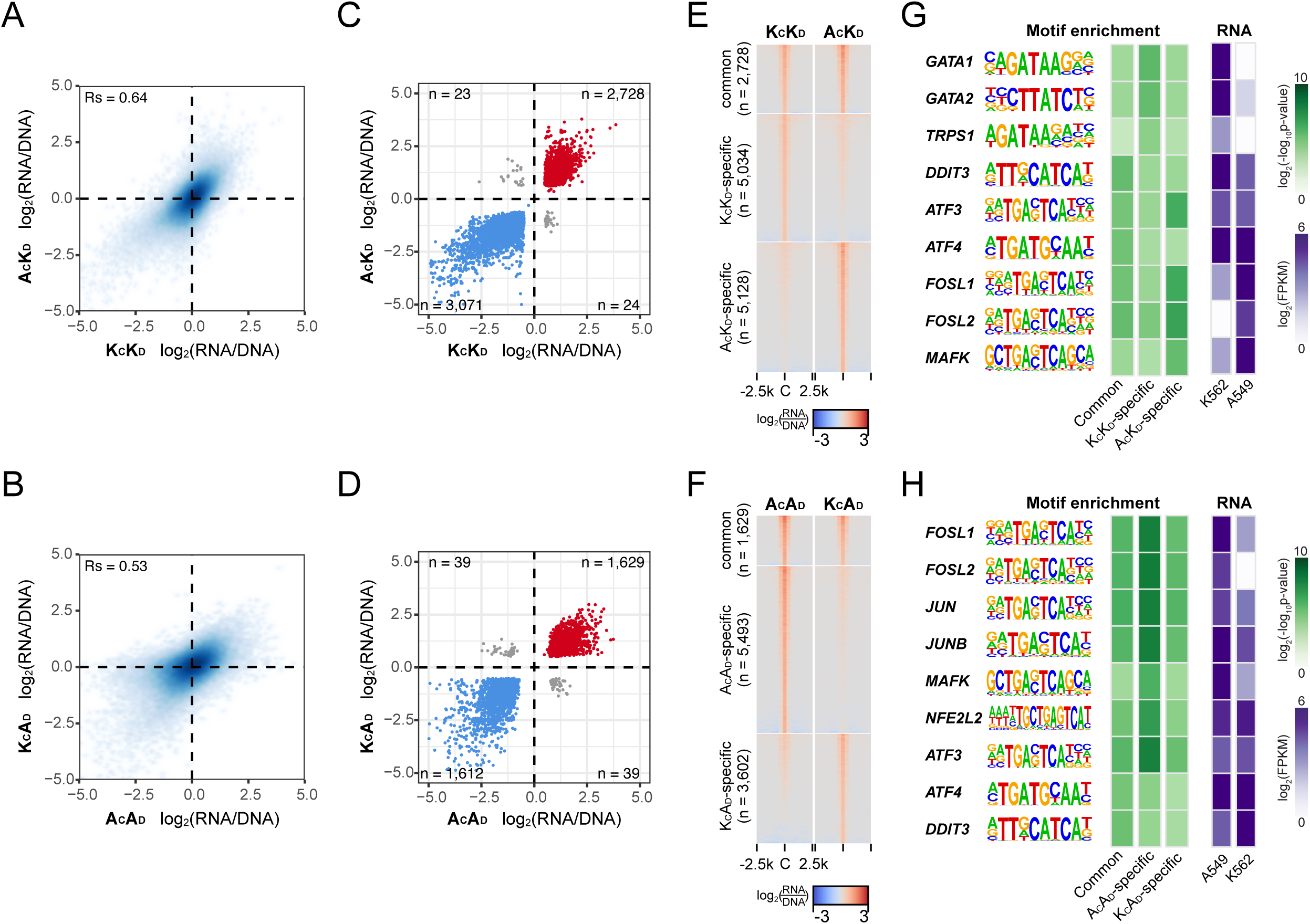
Comparisons between enhancer activities in *cis* and *trans*. (A) Activities comparison of all overlapped sequences when K562 DNA was transfected into K562 cells and A549 cells. Rs: Spearman correlation. (B) Activities comparison of all overlapped sequences when A549 DNA was transfected into A549 cells and K562 cells. Rs: Spearman correlation. (C) Activities comparison of the overlapped significant active enhancers and overlapped significant negative sequences when K562 DNA was transfected into K562 cells and A549 cells. (D) Activities comparison of the overlapped significant active enhancers and overlapped significant negative sequences when A549 DNA was transfected into A549 cells and K562 cells. (E) Heatmap showing the common and condition-specific active enhancers when K562 DNA was transfected into K562 cells and A549 cells. (F) Heatmap showing the common and condition-specific active enhancers when A549 DNA was transfected into A549 cells and K562 cells. (G) The enriched TF motifs for the common and condition-specific active enhancers when K562 DNA was transfected into K562 cells and A549 cells. (H) The enriched TF motifs for the common and condition-specific active enhancers when A549 DNA was transfected into A549 cells and K562 cells.

We analyzed the TF motifs enrichment for the common active enhancers in both *cis* and *trans*, and those only highly active in *cis* or *trans*. For K562 DNA, motifs of *GATA1/2* and *TRPS1* showed higher enrichment in *cis*, and these factors were only highly expressed in K562 cells (Fig. 3G). Motifs of *ATF3/4*, *DDIT3* were enriched in all the conditions, and they were highly expressed in both K562 and A549 cells (Fig. 3G,H). For A549 DNA, motifs of AP-1 transcription factor subunits including *FOSL1/2* and *JUN/JUNB* showed higher enrichment in A549 cells, and these factors were highly expressed in A549 cells (Fig. 3H). These results suggested the DNA sequence motifs and the differentially expressed TFs may contribute to the preferred high activity in *cis* or *trans* condition.

### Deep learning model predicts cell type-specific enhancers from sequences

To investigate whether the cell type-specific active enhancers can be predicted only using the DNA sequences by *in silico* model, we built and trained a convolutional neural network (CNN) deep learning model based on the DeepSTARR (de Almeida et al. 2022) for enhancers in K562 and enhancers in A549 separately (see Methods). The model only took DNA sequences with 350bp in length as input and predicted the probability of being active enhancer in the specific cell line (Fig. 4A). Independent test using the receiver operating characteristic (ROC) curve with leave-chromosome-out showed the model for K562 achieved an area under the curve (AUC) of 0.79 and the model for A549 achieved an AUC of 0.76 (Fig. 4B), which was in line with existing studies showing that CNN-based deep learning models could predict enhancer activities and TF binding with high performance (Wang et al. 2018a; Avsec et al. 2021; Novakovsky et al. 2023).

**Figure 4.**
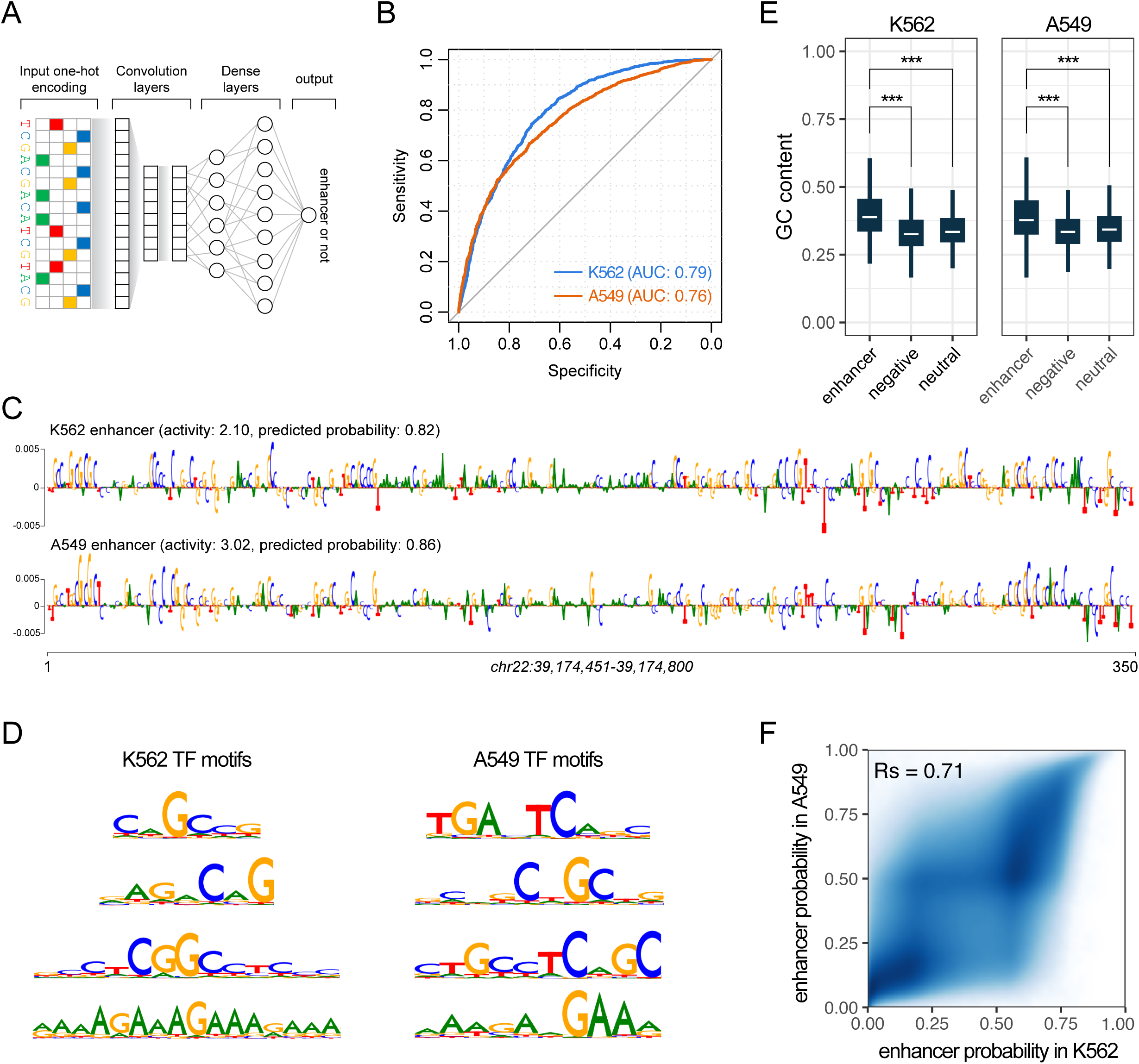
*In silico* prediction of cell type-specific active enhancers from DNA sequences. (A) The architecture of the convolutional neural network (CNN) based deep learning model. (B) Independent test results using the receiver operating characteristic (ROC) curve with leave-chromosome-out. AUC: area under the curve. (C) Example showing the DeepLIFT generated contribution of each nucleotide for the model prediction on the DNA sequence which was an enhancer in both K562 and A549 cells. (D) Cell type-specific high contribution sequence patterns generated by TF-MoDISco. (E) Boxplot showing the GC content distribution among active enhancer sequences, negative sequences and neutral sequences in K562 and A549. P-values were calculated using two-tailed Student’s t-test. ***: p-value<0.001. (F) Comparison of the predicted enhancer probabilities in K562 and A549 for all the whole genome 350bp nonoverlap bins. Rs: Spearman correlation.

One advantage of the CNN-based deep learning models is that the features learned by the CNN reflect potential TF binding preferences, which provides an explanation for the importance and contribution of each nucleotide in the input sequence (Shrikumar et al. 2017; Novakovsky et al. 2023). Figure 4C provided an example enhancer sequence that was active in both K562 and A549 cells. However, the contribution of each nucleotide to the enhancer activity was different in the two types of cells. By summarizing the common high contributing sequence patterns in each model using TF-MoDISco, we found AP-1 motif in A549 model and GAAA motif with high contributions in both models (Fig. 4D). We noticed high G/C number in the high contribution sequence patterns in both models (Fig. 4D). Indeed, when comparing the GC content among the sequences of active enhancers, negative sequences and neutral sequences, the active enhancer sequences showed significantly higher GC content than non-enhancer sequences in both K562 and A549 cells (Fig. 4E), indicating higher GC content would contribute to enhancer activities.

Next, leveraging the deep learning models, we performed *in silico* whole-genome cell type-specific enhancer screen. We split the human GRCh38 reference genome into 350bp nonoverlap bins and predicted their enhancer probabilities in K562 and A549 cells. Comparison of the *in silico* enhancer prediction results in the two cell lines showed high consistency across the whole genome with Spearman correlation coefficient 0.71 (Fig. 4F), which further supported our above finding that the regulatory property of the same sequence kept similar in different types of cells.

## Discussion

In this study, by transfecting and cross-transfecting each enhancers ChIP-STARR-seq library to the native cells and a different type of cells, we generated two genome-wide enhancer *cis* activity maps and two enhancer *trans* activity maps in two different cell lines. Our strategy of specifically enriching and selecting H3K4me1 marked dinucleosomes together with the linker sequence between them enabled us to study enhancer activities in high-resolution. We used the STARR-seq vector with the origin of replication (ORI) as the core promoter which is more suitable for human cells, avoiding potential competition between ORI and other core promoters (Lemp et al. 2012; Muerdter et al. 2018). We not only sequenced the reporter RNA but also the transfected plasmid DNA from cells and used it as the normalization basis in a model-based quantification method (Hansen and Hodges 2022), which allowed us to obtain reliable enhancer activities quantification from two independent replicates in each condition.

The enhancer *cis* activity maps in K562 and A549 cells showed most of the active enhancers were cell type-specific, with different TFs motif enrichment corresponding to the TFs differential expression in the two cell lines. In native chromatin environment, the enhancer activity landscape is shaped by sequence intrinsic features, chromatin state, chromatin interaction and TFs expressed in the cells (Gasperini et al. 2020). Our plasmid-based enhancer activity assay only measured the naked DNA intrinsic contribution under the cell type-specific TFs environment. Due to the enhancer-promoter interaction specificity (Zabidi et al. 2015; Martinez-Ara et al. 2022), the measured enhancer activities in STARR-seq may change when using a different core promoter. Nevertheless, we found most of the active enhancers from STARR-seq displayed active chromatin markers in their native chromosome loci including H3K27ac enrichment, opening chromatin and p300 binding, indicating they were also active in the native chromatin state.

By investigating how the activity landscape of enhancer repository would change when transferred from one type of cells to another type of cells, we found the regulatory property generally kept similar between *cis* and *trans* conditions, suggesting it is the sequence intrinsic feature and independent of TFs environment. However, the levels of activity are quite different for the same sequence in different type of cells, which may be due to the differentially expressed TFs. Altogether, our study provides a new perspective of sequence intrinsic enhancer activities in different types of cells.

## Methods

### Cell culture

Both K562 and A549 cell lines were cultured with 1640 RPMI medium with 10% FBS, Pen/Strep (Gibco 15140-163, 500μl per 50ml) and L-Glutamine (Gibco 25030-164, 500μl per 50ml). The cells were incubated at 37℃ with 5% CO_2_. The K562 cells were split into 0.4×10^6^ cells/ml when the density reached 0.8×10^6^ cells/ml. The A549 cells were split 1:4 when growing into 90% confluent. Each cell type had two independent cell cultures.

### Cell line verification

When the cells growing into log phase, 1×10^6^ cells were harvested from each culture and DNA was extracted using QIAamp DNA mini Kit (QIAGEN 51304). STR analysis was performed with PowerPlex 21 System (Promega DC8902) following the manufactory’s protocol. The genotypes of the STR loci were compared with the records in ATCC STR profile database and each cell line identity was confirmed with 100% match. Free of mycoplasma in each culture was confirmed using PCR-based method with MGSO/GPO-3 primer pairs as described previously (Young et al. 2010).

### Cell sample preparation

After the total number of cells reached 1×10^7^ in each culture, cells were harvested. 1×10^7^ K562 cells were harvested by centrifuge at 300g for 5min and washed 3 times with chilled PBS. 1×10^7^ A549 cells were treated with 1% trypsin for 5min then centrifuged at 300g for 5min and washed 3 times with chilled PBS. Cell pellet was kept and the supernatant was totally removed. Each cell pellet was flash freezed by putting in liquid nitrogen for 2min then stored at -80℃.

### Incomplete MNase digest of chromatin

All the procedures were carried on ice. The flash freezed cells of K562 (2 replicates) and A549 (2 replicates) were put on ice for 15min then resuspended in 250μl chilled douncing buffer (10mM Tris-HCl pH7.5, 4mM MgCl_2_, 1mM CaCl_2_) with 1X Protease Inhibitory Cocktail (PIC, Roche 5892791001). Cells were homogenized through a 1ml 25g 5/8 syringe for 20 times and kept on ice. To digest the chromatin, 0.5ul of 2,000U/μl MNase (NEB M0247S) were added in each homogenate then incubated (no prewarming) at 37℃ for 7min. The reaction was terminated by putting on ice immediately, adding 5μl of 0.5M EDTA and incubated on ice for 5min. Then 1ml fresh prepared chilled hypotonic buffer (0.2mM EDTA pH8.0, 0.1mM benzamidine, 0.1mM phenylmethylsulfonyl fluoride, 1.5mM dithiothreitol) with 1X PIC was added in each reaction and incubated on ice for 1h, with brief vortexing every 20min. The cell debris was pelleted by centrifuging at 4℃, 3,000g for 5min. The supernatant of each sample was transferred to a new 1.5ml LoBind tube and kept on ice. The incomplete digestion was assayed by 1% agarose gel electrophoresis with 30μl supernatant of each sample.

### ChIP of H3K4me1 nucleosome-DNA complex

The MNase digested chromatin solution of each sample was pre-cleared with 100μl Dynabeads Protein A/G 1:1 mix magnetic beads (Invitrogen 10001D, 10003D) and rotated at 4℃ for 2h. Then each tube was put on DynaMag-2 magnet stand to separate the beads from the supernatant and transferring the supernatant to a new 1.5ml LoBind tube. Chromatin immunoprecipitation was carried out by adding 12μl (1μg/μl) anti-histone H3 monomethyl K4 (H3K4me1) antibody (Abcam ab8895) to each tube and rotated at 4℃ for 1h. For the control sample, 10μg rabbit IgG antibody (Sigma I8140) was added and rotated at 4℃ for 1h. The antibody-nucleosome-DNA complex was precipitated by adding 60μl Protein A/G mix magnetic beads and rotated at 4℃ overnight (12h). Beads were recovered by putting on magnet stand and the supernatant was discarded. The beads were washed twice with 500μl ChIP wash buffer I (20mM Tris-HCl pH8.0, 0.1% SDS, 1% Triton X-100, 2mM EDTA, 150mM NaCl) with 1X PIC, followed by a single wash with ChIP wash buffer II (20mM Tris-HCl pH8.0, 0.1% SDS, 1% Triton X-100, 2mM EDTA, 500mM NaCl) with 1X PIC. The beads were eluted in 200μl elution buffer (100mM NaHCO_3_, 1% SDS) with 1μl RNase A, and incubated at 68℃ for 2h. Then each tube was put on the magnet stand and the supernatant was transferred to a new 1.5ml LoBind tube. Elution was repeated with addition of 100μl elution buffer and incubated at 68℃ for 30min. The two elutes of a sample were pooled together. The immunoprecipitated material of each sample was purified using QIAGEN MinElute PCR purification kit (QIAGEN 28004) and the DNA was recovered with 40μl elution buffer (10mM Tris-Cl pH8.5).

### Size selection of dinucleosome DNA

Dinucleosome ChIP DNA fragments were selected using Pippin Prep with 1.5% agarose gel cassette and dye-free internal marker K (Sage Science CDF1510). 30μl of purified immunoprecipitated DNA of each sample was mixed with 10μl marker mix and loaded into each sample well. The collection mode was set to range from 260bp to 490bp. After the run finished, 50μl of size selected DNA in each elution module was transferred to a new LoBind tube. The size selected DNA samples were analyzed using Agilent 2100 Bioanalyzer following the manufactory’s protocol and DNA size was confirmed as expected.

### Library insert generation

50μl of size selected DNA was used as the start material for library generation using NEBNext Ultra II DNA library preparation kit for Illumina (NEB E7645S). The end preparation (end repair and A tailing) and adaptor ligation steps were performed following the manufactory’s protocol. The adaptor-ligated DNA was purified using AMPure XP Beads (Agencourt A63881) following protocol of the Ultra II library prep kit and the purified DNA was recovered with 30μl 10mM Tris-Cl (pH8.5). To amplify the adaptor-ligated DNA, 4 PCR reactions were set up for one sample with 7.5μl purified adaptor-ligated DNA, 7.5μl ddH_2_O, 5μl 10uM forward primer (TAGAGCATGCACCGGACACTCTTTCCCTACACGACGCTCTTCCGATCT), 5ul 10μM reverse primer (GGCCGAATTCGTCGAGTGACTGGAGTTCAGACGTGTGCTCTTCCGATCT), and 25μl 2X Kapa HiFi HotStart ReadyMix (Kapa KK2611). The PCR program was 98℃ for 45s, 10 cycles of 98℃ - 15s, 65℃ - 30s and 72℃ - 45s, final elongation at 72℃ for 60s and hold at 4℃. The PCR products for one sample were pooled together and purified with AMPure XP beads, using 0.9 volume beads per 1 volume PCR reaction and eluted with 200μl 10mM Tris-Cl (pH8.5). To increase cloning efficiency, the amplified library DNA was further purified using QIAGEN MinElute PCR purification kit (QIAGEN 28004) and the DNA was eluted with 50μl elution buffe.

### STARR-seq plasmid library preparation

The STARR-seq vectors (Addgene #99296) were linearized by restriction enzyme digestion using 2μl AgeI-HF (NEB R3552S) and 2μl SalI-HF (NEB R3138T) at 37℃ overnight. The digested vectors were run on 1% agarose gel and the target band was cut and harvested using QIAquick Gel Extraction Kit (QIAGEN 12362). The gel extraction products were further purified with QIAGEN MinElute PCR purification kit (QIAGEN 28004). The library inserts from last step were ligated to the linearized STARR-seq vectors using In-Fusion HD Cloning Kit (Clontech 639650) following the manufactory’s protocol. For each library, 10 In-Fusion cloning reactions were performed and every 5 reactions were pooled together. Each product was adjusted to 250μl by adding Elution Buffer. Then 25μl 3M NaAc (pH 5.2), 2μl Pellet Paint NF Co-Precipitant (Millipore 70748-3) and 750μl ice-cold 100% EtOH were added and put at -20℃ for 16h to precipitate the DNA. The DNA pellet was washed twice with ice-cold 70% EtOH, centrifuged at full speed for 15min at 4℃ and air dried at room temperature. The pellet was resuspended with 12μl Elution Buffer and stored at -20℃. To clone the plasmid library, 2μl of the purified In-Fusion cloning products were transfected into 25μl E. cloni 10G ELITE Electrocompetent Cells (Lucigen 60052-4) in 1mm cuvette (Bio-Rad 1652089) using Bio-Rad Gene Pulser Xcell electroporation with 1800v, 10μF and 600Ω. For each library, 10 reactions were performed and pooled together then cultured in 3L LB+Amp medium at 37℃ 300rpm for 15h. The plasmid library was extracted using QIAGEN Plasmid Plus Maxi Kit (QIAGEN 12963). The product was further purified using ethanol precipitation and resuspended in pure H_2_O. The final concentration of each plasmid library was adjusted to 1μg/μl.

### STARR-seq screening in K562 and A549 cells

The STARR-seq screening was performed as described previously (Muerdter et al. 2018). Each plasmid library was transfected into K562 or A549 cells by electroporation using Lonza SF Cell Line 4D-Nucleofector X Kit (Lonza V4XC-2012) with Lonza 4D-Nucleofector. For each library, 20 electroporation reactions were performed with 5×10^6^ cells and 10μg plasmid library per reaction, following the manufactory’s protocol. After transfection, the cells were cultured for 24h at 37℃ in culture mediums without Pen/Strep. Then the cells were harvested and 3/4 of the cells were used for RNA extraction while the other 1/4 were used for plasmid DNA extraction. The RNAs were extracted using QIAGEN RNeasy Midi Kit (QIAGEN 75144) and mRNAs were selected with Invitrogen Dynabeads mRNA Purification Kit (Invitrogen 61006) following the manufactory’s protocol. The mRNAs were treated with Turbo DNase I (Ambion AM2238) and cleaned up using QIAGEN RNeasy MinElute clean up kit (QIAGEN 74204) and eluted in 60μl RNase-free water. Reverse transcription of the mRNAs was performed with 59μl eluted mRNA, 5μl dNTP and 1μl 10μM reporter gene specific primer: CTCATCAATGTATCTTATCATGTCTG using the Invitrogen SuperScript III First-Strand Synthesis System (Invitrogen 18080051) following the manufactory’s protocol. The cDNA was treated with RNase A and cleaned up with AMPure XP beads (Beckman A63882). Junction PCR was performed to enrich the reporter cDNA with forward primer TCGTGAGGCACTGGGCAG*G*T*G*T*C, reverse primer CTTATCATGTCTGCTCGA*A*G*C and 2X Kapa HiFi HotStart ReadyMix (Kapa KK2611). The sequencing ready library was generated by PCR with the junction PCR product, Universal PCR Primer for Illumina and index primer for Illumina (NEB E7335S), and 2X Kapa HiFi HotStart ReadyMix (Kapa KK2611). The PCR product was purified using SPRIselect beads (Beckman B23318). The transfected plasmid DNA was extracted from the cells using QIAGEN Plasmid Plus Midi kit (QIAGEN 12943). The sequencing ready plasmid library was generated by PCR with the extracted plasmid, Universal PCR Primer for Illumina and index primer for Illumina (NEB E7335S), and 2X Kapa HiFi HotStart ReadyMix (Kapa KK2611). The PCR product was purified using SPRIselect beads (Beckman B23318).

### STARR-seq data analysis

The raw sequencing data for reporter RNAs and plasmid DNAs were trimmed with Trimmomatic (version v0.39) (Bolger et al. 2014) to remove adaptors. The cleaned reads were mapped to human reference genome GRCh38 using bwa mem (version 0.7.17) (Li and Durbin 2010). The mapped reads were filtered with minimum mapping quality of 30 and minimum fragment length of 250. Potential PCR duplicates were kept. The signal tracks were generated with deeptools bamCoverage (version 3.5.2) (Ramirez et al. 2016).

### Active enhancer analysis

To calculate enhancer activities, we split the human GRCh38 reference genome into 350bp nonoverlap bins and counted the number of RNA fragments and DNA fragments in each bin for each replicate with bedtools (version 2.31.0) (Quinlan and Hall 2010). We filtered the bins with DNA fragments count per million (CPM) less than 1 in either replicate. Then we calculated the p-value and fold change of RNA over DNA using DESeq2 (version 1.38.3) (Love et al. 2014) by leveraging the two replicates of RNA and DNA and using the DNA count as control. For DESeq2, local fit type was used. Bins with raw p-value < 0.05 and log2 fold change > 0 were considered as active enhancers. Bins with p-value < 0.05 and log2 fold change < 0 were considered as negative sequences. Bins with p-value > 0.95 and the absolute value of log2 fold change < log_2_(1.1) were considered as neutral sequences.

The distribution of enhancers was analyzed with ChIPseeker (version 1.34.1) (Yu et al. 2015). GO enrichment was performed with clusterProfiler (version 4.6.2) (Yu et al. 2012). Deeptools (version 3.5.2) was used to plot signal enrichment heatmaps. Motif enrichment was calculated using HOMER (version 4.11) (Heinz et al. 2010). The transposon annotations for human GRCh38 genome were downloaded from RepeatMasker (https://www.repeatmasker.org/species/hg.html).

### Replicates correlation

For replicates correlation evaluation of ChIP-seq and STARR-seq plasmid DNA, we used 1kb bin size across the genome and calculated the Pearson correlation. For replicates correlation of STARR-seq RNA, we used 350bp bin size and only considered the bins with corresponding DNA fragment count per million ≥1 in both replicates then calculated the Pearson correlation.

### Public data usage

The following data in K562 cells from ENCODE project (Encode Project Consortium et al. 2020) were used: RNA-seq (ENCFF421TJX), H3K27ac ChIP-seq (ENCFF381NDD), ATAC-seq (ENCFF102ARJ), p300 ChIP-seq (ENCFF636VVR), ChromHMM annotation (ENCFF963KIA). The following data in A549 cells from ENCODE project were used: RNA-seq (ENCFF024DUU), H3K27ac ChIP-seq (ENCFF105IIK), ATAC-seq (ENCFF872SDF), p300 ChIP-seq (ENCFF134ILE), ChromHMM annotation (ENCFF418WHV).

### Deep learning model for enhancer prediction

The deep learning model was built based on DeepSTARR (de Almeida et al. 2022). The model used 350bp DNA sequences as input and converted the sequence to one-hot representation. We used three convolution layers with number of filters 64, 16, 16 for each layer and kernel sizes of 7, 5, 5 for each layer. Then two dense layers were connected with number of neurons 32 and 64 for each layer. The model predicted the enhancer probability of the input sequence. The model was implemented with Keras and TensorFlow (version 2.13.0). We generated independent testing data with enhancers and non-enhancers (negative sequences and neutral sequences) on chromosome 19-22. Validation data were composed of enhancers and non-enhancers on chromosome 16-18. Enhancers and non-enhancers on the other chromosomes were used as training data. The model was trained with Adam as the optimizer and binary cross-entropy as the loss function. The ROC analysis was performed with R package pROC (version 1.18.4). The contribution score of each nucleotide was calculated using DeepLIFT (Shrikumar et al. 2017) and Shap (version 0.42.1). The common patterns of features were identified with TF-MoDISco (version 0.5.16).

## Data access

All raw and processed sequencing data generated in this study have been submitted to the NCBI Gene Expression Omnibus (GEO; https://www.ncbi.nlm.nih.gov/geo/) under accession number GSE243793. The code used in this study is available at https://github.com/icebert/ChIP-STARR-seq.

## Acknowledgement

This work is impossible without the kind help from many persons. We are grateful to Dr. Liping Wei for her great help in this project. We thank Dr. Alexander Stark for kindly providing the STARR-seq vectors. We thank Dr. Tenghui Yu at Peking University for his great help on cells electroporation. We thank Dr. Ge Gao, Dr. Lichen Ren and Dr. Cheng Li at Peking University for their help on cell culture. We are grateful to Hongxia Lv and Liying Du from the core facility at National Center for Protein Sciences at Peking University, for their help on electroporation and flow cytometry. We acknowledge Dr. Junlin Teng at Peking University for her help on bacteria electroporation. This work was supported by the National Natural Science Foundation of China (No. 31530092) and the Ministry of Science and Technology of China (No. 2015AA020108). M.W. was supported in part by the Postdoctoral Fellowship of Peking-Tsinghua Center for Life Sciences.

## Author contributions

M.W. and Q.W. conceived the project. M.W. performed the experiments with help from X.Y. and Q.W. M.W. performed the data analysis. M.W. wrote the manuscript. All authors reviewed the manuscript.

