## Supplemental Figures for "High-resolution dissection of human cell type-specific enhancers in *cis* and *trans* activities"

### Supplemental Figure S1

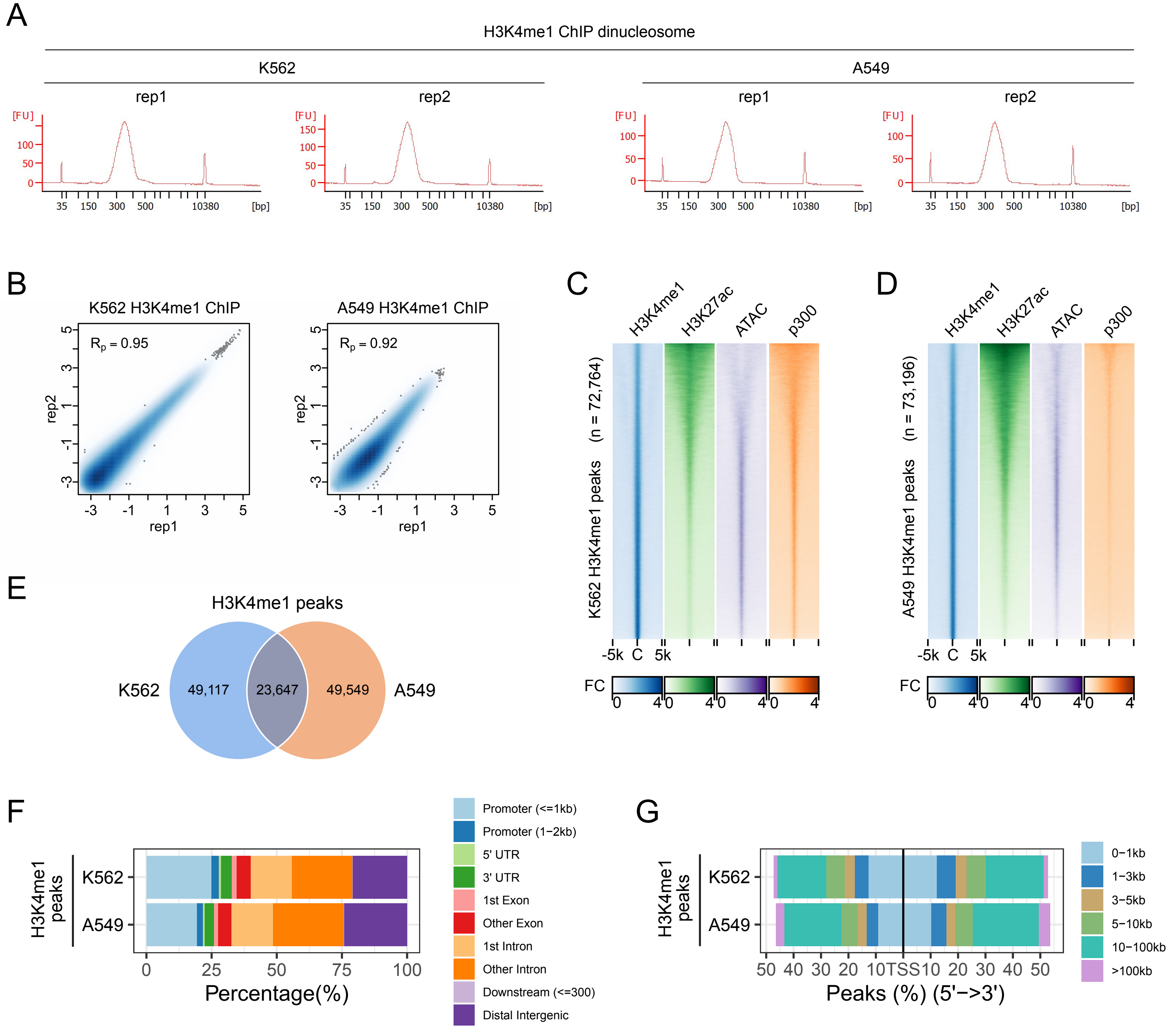

**Supplemental Figure S1. Evaluation of the H3K4me1 dinucleosome ChIP sequences with ChIP-seq.** (A) H3K4me1 ChIP dinucleosome DNA fragment size evaluation using the Agilent 2100 Bioanalyzer. (B) ChIP-seq replicates correlation for K562 and A549. Rp: Pearson correlation. (C) Heatmaps showing the enrichment of H3K27ac, ATAC-seq and p300 binding signals for the K562 H3K4me1 peak regions. (D) Heatmaps showing the enrichment of H3K27ac, ATAC-seq and p300 binding signals for the A549 H3K4me1 peak regions. (E) Overlaps of H3K4me1 peaks between K562 and A549 cells. (F) Genomic distribution of H3K4me1 peaks in K562 and A549. (G) Distance distribution of H3K4me1 peaks to their nearest transcription start site (TSS) in K562 and A549.

### Supplemental Figure S2

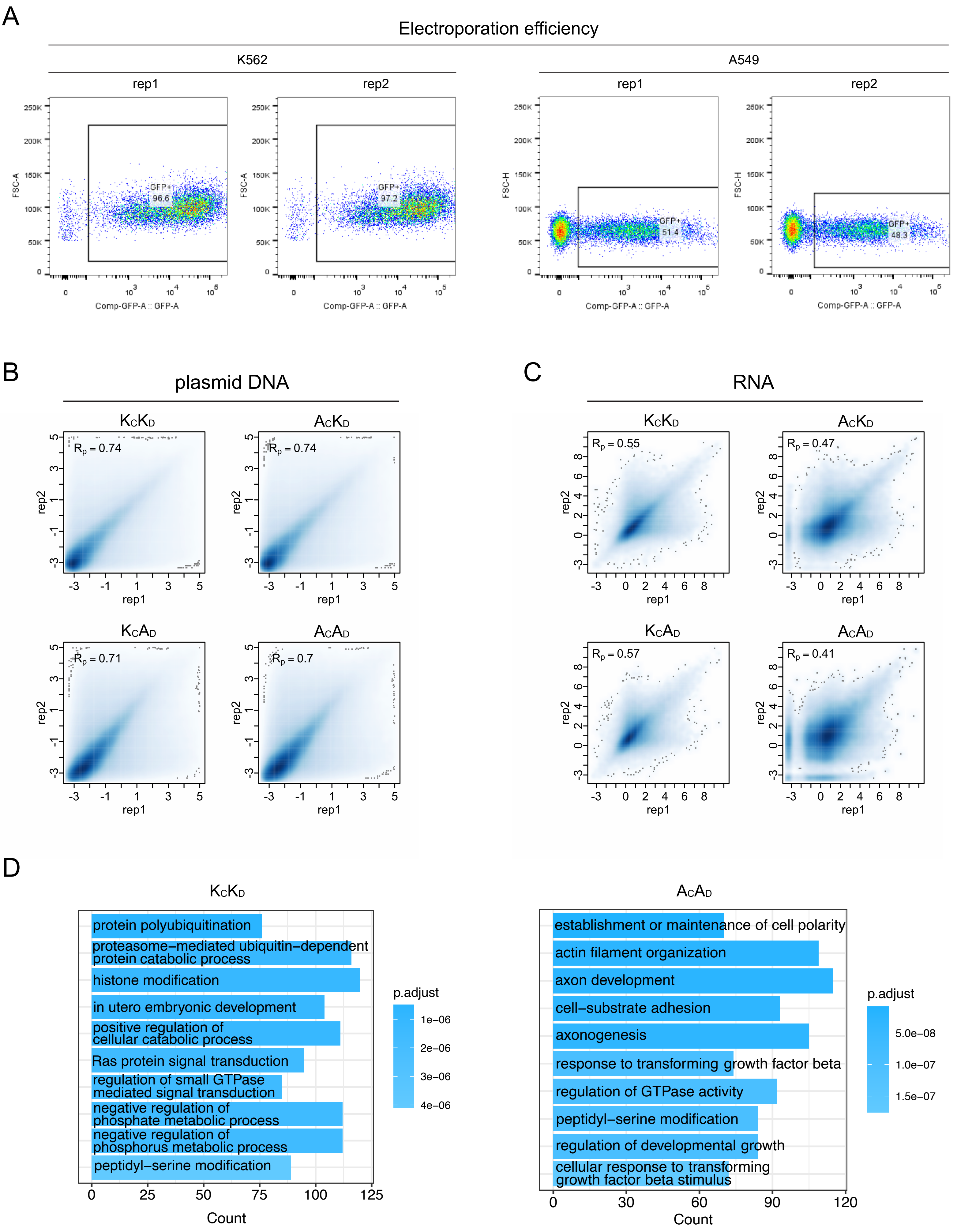

**Supplemental Figure S2. Evaluation of the ChIP-STARR-seq data quality.** (A) Percentage of GFP+ cells after electroporation for K562 cells and A549 cells, measured by flow cytometry. For each type of cell, two replicates were performed. (B) Replicates correlation for the isolated plasmid DNA from cells after transfection. Rp: Pearson correlation. $K_CK_D$ – K562 cells with K562 DNA; $A_CA_D$ – A549 cells with A549 DNA; $A_CK_D$ – A549 cells with K562 DNA; $K_CA_D$ – K562 cells with A549 DNA. (C) Replicates correlation for the reporter RNAs from cells after transfection. Rp: Pearson correlation. (D) Gene Ontology (GO) terms enrichment for the closest genes to the active enhancers in K562 (left) and in A549 (right).

### Supplemental Figure S3

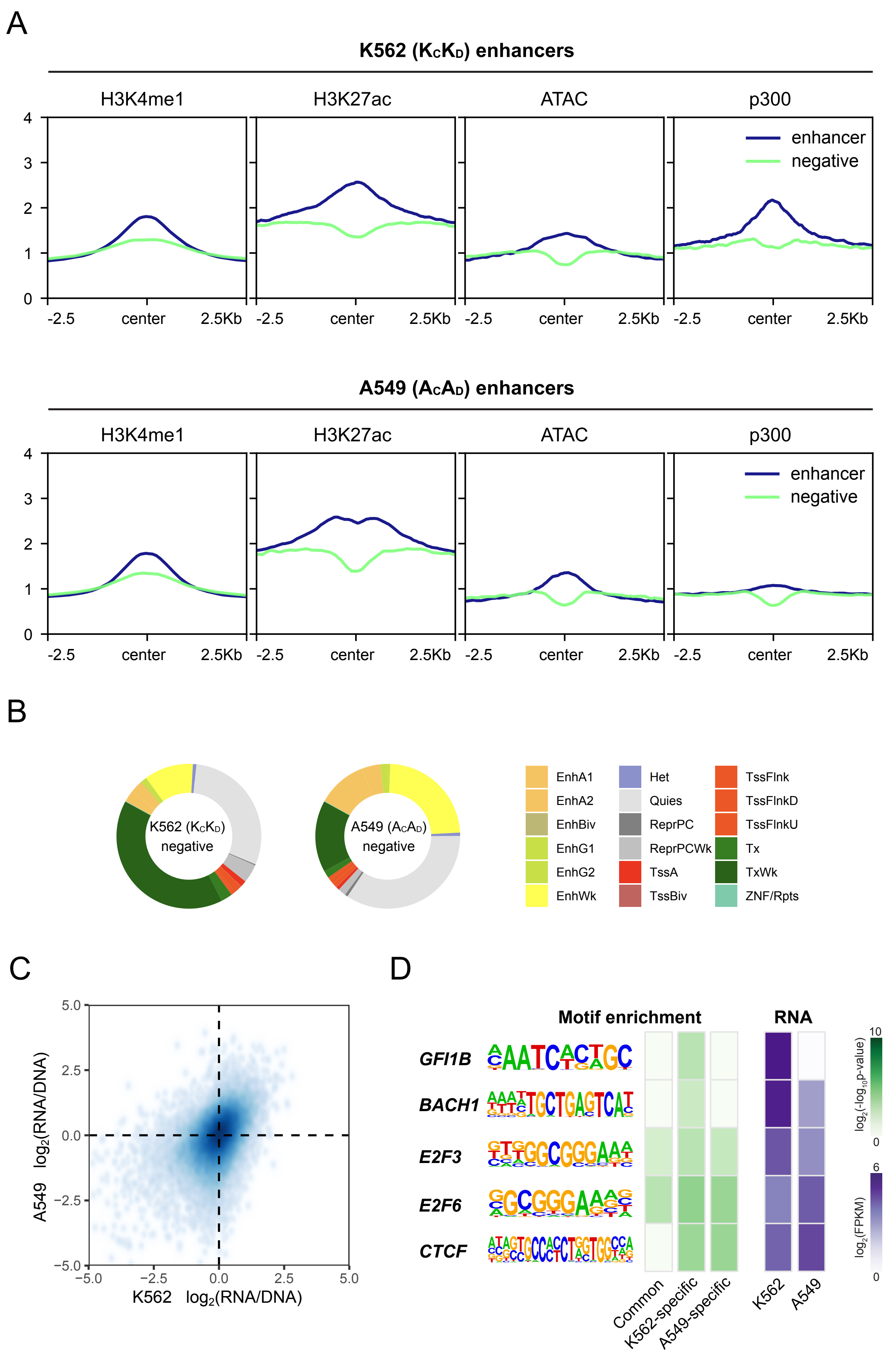

**Supplemental Figure S3. Cell-type specific enhancers and negative sequences.** (A) Profile of H3K4me1, H3K27ac, chromatin accessibility (measured by ATAC-seq) and p300 binding signal at the genomic regions of active enhancers compared to negative sequences in K562 cells and in A549 cells. (B) Overlaps of negative sequences with ChromHMM annotation. EnhA1 - Active enhancer 1; EnhA2 - Active enhancer 2; EnhBiv - Bivalent enhancer; EnhG1 - Genic enhancer 1; EnhG2 - Genic enhancer 2; EnhWk - Weak enhancer; Het - Heterochromatin; Quies - Quiescent/low; ReprPC - Repressed Polycomb; ReprPCWk - Weak repressed Polycomb; TssA - Active TSS; TssBiv - Bivalent/poised TSS; TssFlnk - Flanking TSS; TssFlnkD - Flanking TSS downstream; TssFlnkU - Flanking TSS upstream; Tx - Strong transcription; TxWk - Weak transcription; ZNF/Rpts - ZNF genes & repeats. (C) Activities comparison of all overlapped sequences between K562 and A549. (D) For negative sequences, the enriched motifs of TFs in each group and the expression levels of the TFs in K562 and A549.
